# Multiaxial Biophysical Control of Oncogenic Phase Separation by Indoleamines: A Proof-of-Concept Synthesis of Landscape-Level Regulation

**DOI:** 10.64898/2026.02.03.703596

**Authors:** Doris Loh, Luiz Gustavo de Almeida Chuffa, Fábio Rodrigues Ferreira Seiva, Russel J. Reiter

## Abstract

**Background:** Phase-separated oncogenic condensates act as biophysical sanctuaries, stabilizing transcriptional and survival programs to drive cancer progression. While individual proteins within these assemblies are well-documented, a universal regulator capable of orchestrating the integrated biophysical axes governing cellular phase behavior has remained elusive. We propose a paradigm of landscape-level regulation mediated by the indoleamine scaffold.

**Methods:** A systematic synthesis and integrative bioinformatics analysis were performed to identify the intersection between melatonin-responsive genes and the phase-separation proteome. A gene set of 121 phase separation-related molecules was validated via PhaSepDB and intersected with 134 melatonin-modulated genes identified across diverse cancer models. Clinical relevance was assessed using TCGA survival datasets.

**Results:** We identified a conserved network of 26 core genes—including *AR, BCL2, CGAS, CTNNB1, EP300, EZH2, EGFR, IKBKG (NEMO), KEAP1, KDM1A (LSD1), LEF1, MYC, NANOG, PRNP, SMAD3, SOX9, SQSTM1, TFEB, TFAM, TP53, TWIST1, USP10, WWTR1 (TAZ), VIM, YAP1*, and *YTHDF3*—at the intersection of melatonin signaling and PS architecture. Network analysis revealed high-confidence interactions predominantly localized to the nucleoplasm and stress granules. Melatonin significantly suppressed the majority of these oncogenic drivers, targeting pathways associated with EMT, glycolysis, and hypoxia. Furthermore, higher expression of this melatonin-responsive phase separation-network correlated with poorer overall survival in breast and gastric cancers.

**Conclusions:** This proof-of-concept synthesis positions melatonin as a remarkably versatile regulator of phase-separated oncogenic networks, capable of simultaneously modulating the integrated biophysical axes that govern cellular phase behavior. We propose that melatonin utilizes a tri-lever framework—comprising redox switching, structural intercalation, and dielectric tuning—to recalibrate the cellular solvation and fluidity landscape. By enforcing fundamental physicochemical constraints rather than targeting discrete signaling nodes, melatonin destabilizes the biophysical sanctuaries that shield oncogenic programs. This landscape-level regulation offers a strategic platform for disrupting condensate-driven malignancy, providing a biophysical logic that circumvents the diverse evasion mechanisms of the malignant cell.

## 1 Introduction

Phase separation is an evolutionarily conserved thermodynamic process utilized by all tested living organisms across the three major domains of life to organize and sustain fundamental biological processes. Under physiological conditions, phase separation is a reversible process that forms dynamic, membraneless, micron-scale, liquid-like compartments known as biomolecular condensates (BCs). These condensates can rapidly respond to fluctuating cellular environments. The formation and maintenance of BCs are driven by nonequilibrium thermodynamic conditions arising from the competition between entropy and enthalpy. BCs organize and reorganize cellular biochemistry through the selective inclusion or exclusion of substrates, thereby tuning, promoting, or inhibiting cellular functions and reactions [1–3]. Aberrant phase separation that irreversibly transitions fluid condensates into solid aggregates, resulting in the loss of normal cellular functions, is now a major therapeutic target in oncology [4–7]. In addition to mutations, dysregulation of pH and redox balance in the tumor microenvironment (TME) promotes aberrant phase separation that can inhibit tumor-suppressor genes and enhance oncogene expression [8–10]. Intriguingly, cancer cells often sustain an abnormally reduced intracellular redox state to preserve the liquid-like integrity of redox-sensitive condensates that coordinate core transcriptional and epigenetic programs essential for malignancy [11–13]. Pro-oxidative interventions that selectively perturb these condensates can disrupt their oncogenic functions and relieve repression of tumor suppressor genes [14]. In parallel, recent advances show that phase separation modulates the folding and stability of redox-sensitive, non-canonical DNA structures, including G-quadruplexes and i-motifs, which in turn influence gene activation and repression [15–18]. Together, these insights strengthen the emerging rationale for developing cancer therapeutics that target aberrant phase separation.

The biosynthesis of melatonin (N-acetyl-5-methoxytryptamine) is evolutionarily conserved and has been detected in all tested representatives of Bacteria [19], Eukarya [20], and Archaea [21]. These three domains all originated under a largely anoxic atmosphere during early evolution [22]. The identification of the melatonin biosynthetic pathway in Archaea may suggest that the physiological roles of melatonin in early life forms extended beyond its classic antioxidant and pro-oxidant functions [23,24]. First isolated from the bovine pineal gland in 1958 [25], melatonin is now recognized to be produced predominantly outside the pineal gland, with mitochondrial synthesis in vertebrates estimated to account for more than 95% of total organismal production [26,27]. Since the first in vivo demonstration of melatonin’s anti-tumor effects in 1973 [28], research on melatonin and cancer has expanded substantially across experimental, translational, and clinical domains. Melatonin has been reported to exert multifaceted, both direct and indirect, antitumor effects across a broad spectrum of molecular pathways, including antioxidant and pro-oxidant activities [29,30], apoptosis and autophagy [31,32], angiogenesis and metastasis [33,34], regulation of cell proliferation and cell-cycle arrest [35,36], epigenetic modification [29,37], immunomodulation [38], metabolic reprogramming [39–41], modulation of signaling pathways [42–44], and enhanced sensitization to therapy [45,46]. To date, a universal mechanism that can satisfactorily account for the remarkable versatility of melatonin’s protective actions against tumorigenesis has not been elucidated.

The search for a unifying mechanism necessitates a shift toward the fundamental chemical architecture of the molecule. The indoleamine scaffold, of which melatonin is the prototypical representative, possesses unique biophysical attributes that allow it to interact with a vast array of macromolecular assemblies. The amphiphilic nature and “privileged structure” of the indole[47] are intrinsically linked to its ability to engage in redox, multivalent, and electrostatic interactions [48,49]. By viewing melatonin through the lens of an indoleamine-mediated landscape regulator, we can begin to theorize how it might orchestrate cellular behavior at a scale beyond individual molecular pathways.

Guided by this biophysical perspective, the objective of this integrative systematic review and bioinformatics analysis is to identify genes implicated in human cancer that (1) have experimentally validated roles in biomolecular phase separation and (2) are independently reported to be regulated by melatonin, albeit without prior assessment of their condensate behavior. Building on substantial conceptual and mechanistic evidence—derived from peer-reviewed reviews [50–57] and supported by emerging experimental findings [58]—indicating that this indoleamine can modulate phase-separation dynamics in specific pathological contexts, we investigate whether the intersection of these two gene sets suggests a potential biophysical basis for melatonin-dependent modulation of condensate functions relevant to tumorigenesis. We further evaluate the relevance of these genes to cancer initiation, progression, and metastasis. Collectively, this work aims to define and characterize genes jointly regulated by phase separation and melatonin and to assess the therapeutic potential of this landscape-level modulation in cancer processes. Accordingly, our integrative study addresses the following questions: which cancer-associated genes are jointly linked to biomolecular phase separation and melatonin regulation, and what functional and therapeutic significance does this convergence imply?.

## 2 Methods

This systematic review and bioinformatics analysis was conducted in accordance with the Preferred Reporting Items for Systematic Reviews and Meta-Analyses (PRISMA 2020) guidelines [59].

### 2.1 Data Sources and Search Strategy

A systematic search of PubMed and Web of Science was conducted in April 2025 using predefined search terms (‘melatonin AND cancer’ and ‘phase separation AND cancer’). The initial search was carried out by one author, and all authors reviewed and approved the final set of studies included and excluded from the review

### 2.2 Data Collection, Study Screening, and Selection Criteria

All retrieved records were imported into Rayyan [60], a web-based application that facilitates efficient systematic and other literature reviews, for screening and deduplication. A total of 14,090 references were imported into Rayyan. Rayyan flagged 7,099 potential duplicates; 3,596 duplicate records were deleted prior to screening, 3,492 duplicate clusters were resolved (retaining one record per cluster), and 11 flagged records were judged to be not duplicates. After deletion, 10,494 records remained and proceeded to title/abstract screening.

Studies were excluded if they were: (i) review articles, (ii) clinical trials, (iii) studies involving adjuvants or co-treatments administered in addition to melatonin or (iv) studies with irrelevant topics. Studies were included if they met either of the following criteria: (i) investigation of phase separation of genes in cancer, or (ii) investigation of genes modulated by melatonin in cancer.

After screening and full-text assessment, 462 articles remained for detailed review. Of these, 207 studies met all inclusion criteria and were retained for analysis (Supplementary Material: Tables S1–S2 contain the full set of genes extracted from included studies), while 255 were excluded. No conflicts or unresolved decisions remained following screening. The study selection process is summarized in the PRISMA 2020 flow diagram (Figure 1).

**Figure 1:**
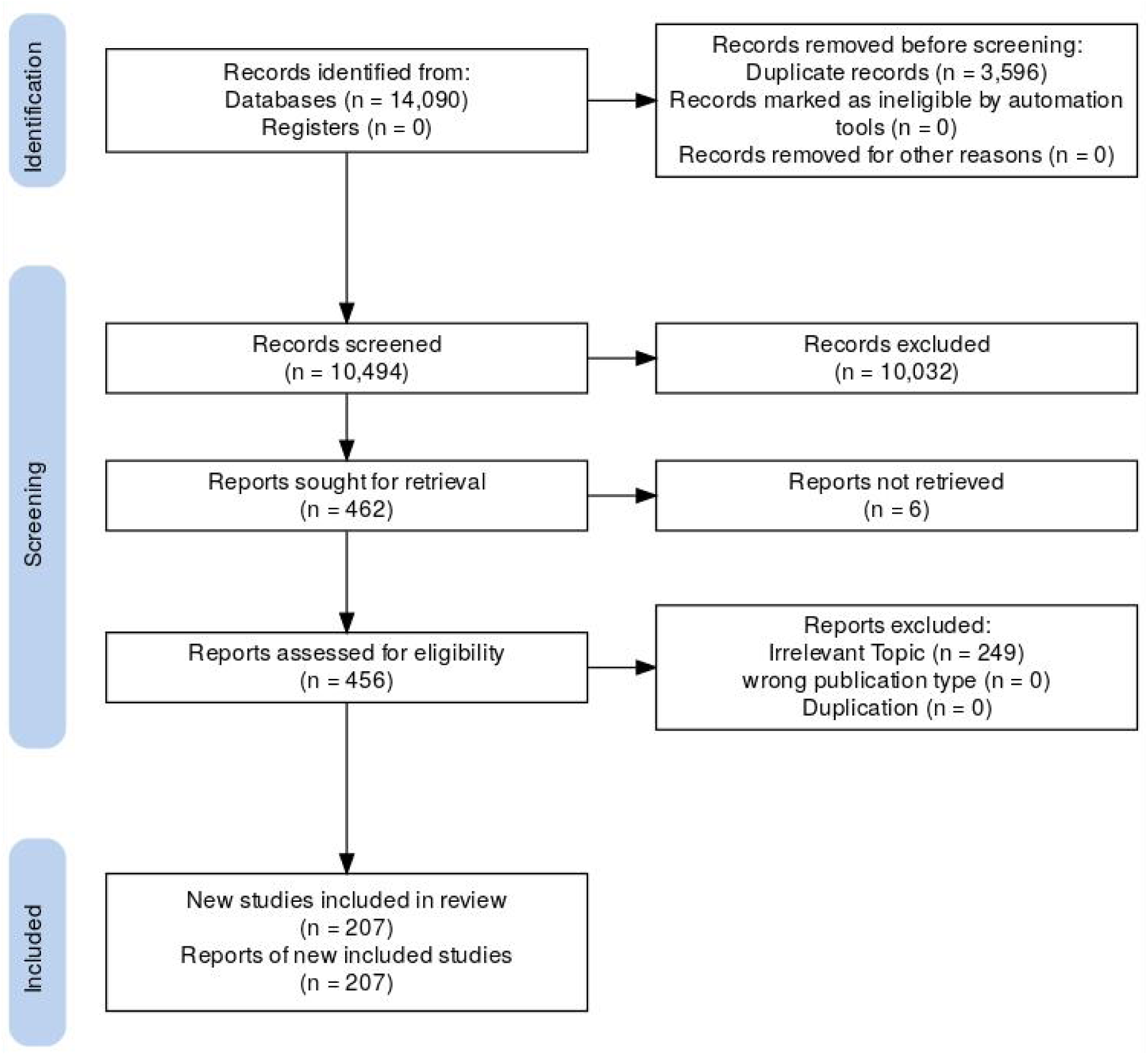
PRISMA 2020 flow diagram for study selection

### 2.3 Bioinformatics Analysis

To check for shared genes between our manual search and those listed in the Phase Separation Database (https://db.phasep.pro/, accessed in October 2025), we used a Venn diagram [61]. Briefly, Phase Separation Database (PSepDB) provides detailed information on proteins and biomolecules involved in liquid-liquid phase separation, helping to advance the understanding of cellular organization and the role of LLPS in various biological processes and diseases [62].

Next, we elaborated a protein-protein interaction (PPI) network employing the STRING database (https://string-db.org/, accessed in October 2025). Using STRING we also consulted the Enriched Pathways and Gene Ontology (GO) terms related with the set of shared genes. The STRING parameters adopted were a complete STRING network with interaction sources from text mining, experimental evidence, databases, co-expression, neighborhood, gene fusion, and co-occurrence, and minimum interaction score of 0.9. STRING database enables the analysis and visualization of protein interaction networks, aiding in the understanding of biological mechanisms underlying various diseases and cellular processes [63]. Cytoscape, version 3.9.1, was used to visualize the PPI and microRNA–gene regulatory networks, and functional co-occurrence.

GO and Pathways enrichment were visualized using ggplot2 R Studio package. Transcription factor (TF) associated with our gene set were explored using the TRRUST v.2 and X2Kweb online databases [64]. The relationships among TF, genes and types of cancer were illustrated through an alluvial diagram, generated with the ggalluvial R Studio package. To search for miRNA-related genes involved in phase-separation, the multMiR R studio package were used. For thus analysis, the parameters were as follow: target = genes, org = “hsa”, table = “validated”, predicted.cutoff.type = “p”, predicted.cutoff = 20, predicted.site = “conserved”, summary = T. Only miRNAs validated by Luciferase Reporter Assay were selected for further analysis. The overall survival data were retrieved from the GEPIA repository (http://gepia2.cancer-pku.cn, accessed in November 2025).

## 3 Results

### 3.1 Melatonin Potentially Alters Phase Separation–Related and Cancer-Associated Genes, Suppressing Major Oncogenic Pathways

After a rigorous screening of the included studies, we identified a total of 121 phase separation-related genes across different biological contexts (Table 1). To better understand the roles of genes associated with phase separation, we first validated this gene set using PhaSepDB, a comprehensive repository providing curated annotations on phase separation-prone molecules including intrinsic determinants, functional relevance, and extrinsic regulatory factors. Of the 121 phase separation-associated genes, 82 were confirmed in PhaSep database (Figure 2A). Pathway enrichment analysis revealed that these genes were predominantly associated with Wnt/β-catenin signaling, Hippo signaling regulation, Extracellular vesicle-mediated signaling in recipient cells (Wiki pathway). Additional enriched pathways included transcriptional regulation by *RUNX3* and *SMAD2/3/4*-mediated transcriptional regulation (Reactome) (Figure 2B). Gene Ontology analysis further demonstrated enrichment of biological processes such as regulation of RNA biosynthesis and DNA-templated transcription, and regulation of epithelial-mesenchymal transition (EMT). The main molecular functions represented included transcription coregulator binding, cis-regulatory region binding, and nuclear receptor binding. These genes were notably localized to key cellular compartments involved in RNA and protein dynamics, including the nucleus, P-bodies, cytoplasmic stress granules, and intracellular membrane-bound organelle (Figure 2C).

**Figure 2:**
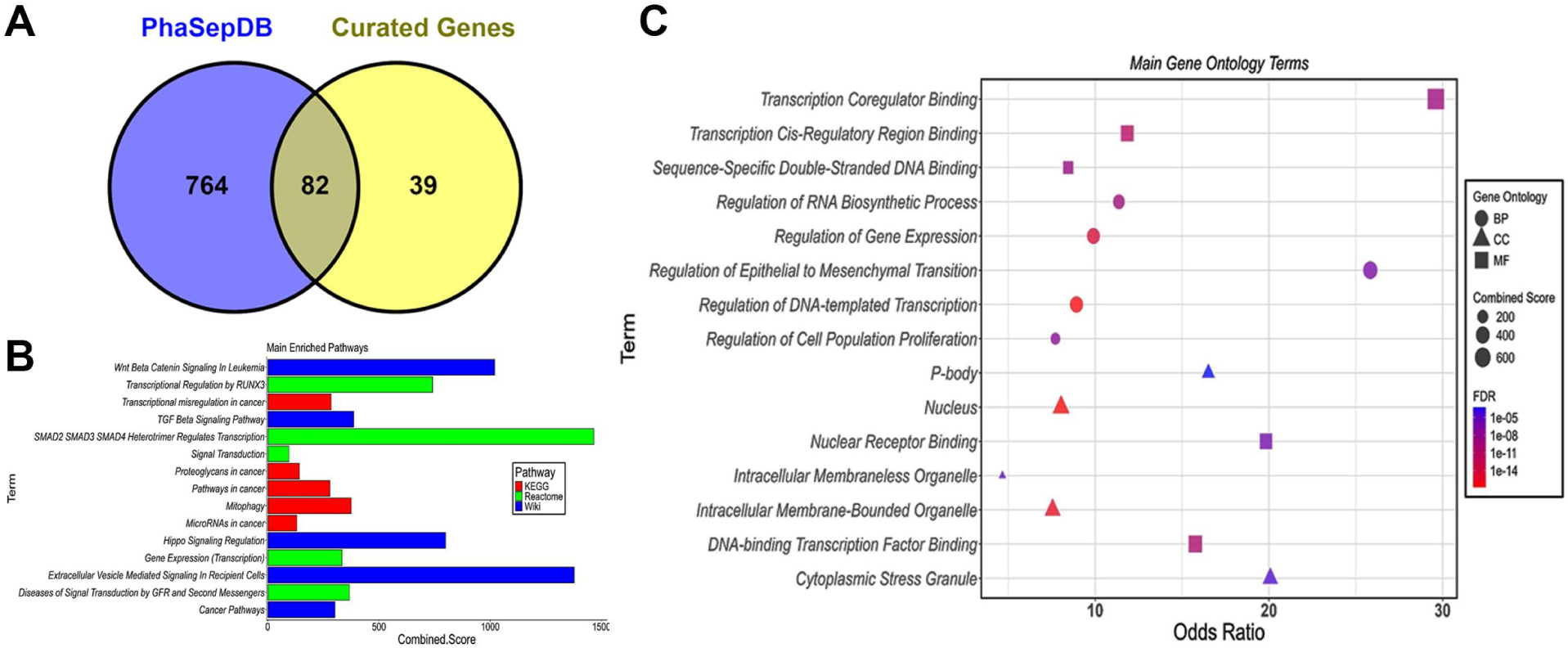
Specific genes involved with phase-separation and functional pathway enrichment. A) Venn diagram showing the overlap between curated genes and PhaSepDB entries. B) Pathway enrichment analysis of the overlapping gene set using KEGG, Reactome, and WikiPathways. Combined scores indicate the strength of pathway enrichment across databases. C) Gene Ontology (GO) enrichment analysis highlighting major biological processes (BP), cellular components (CC), and molecular functions (MF) associated with the intersecting genes. Terms with the highest odds ratios include transcriptional regulation, epithelial-to-mesenchymal transition, RNA biosynthetic processes, DNA-binding transcription factor activity, and localization to intracellular membrane-bounded organelles, P-bodies, and stress granules. Dot size represents combined enrichment score, while color indicates FDR significance.

**Table 1:**
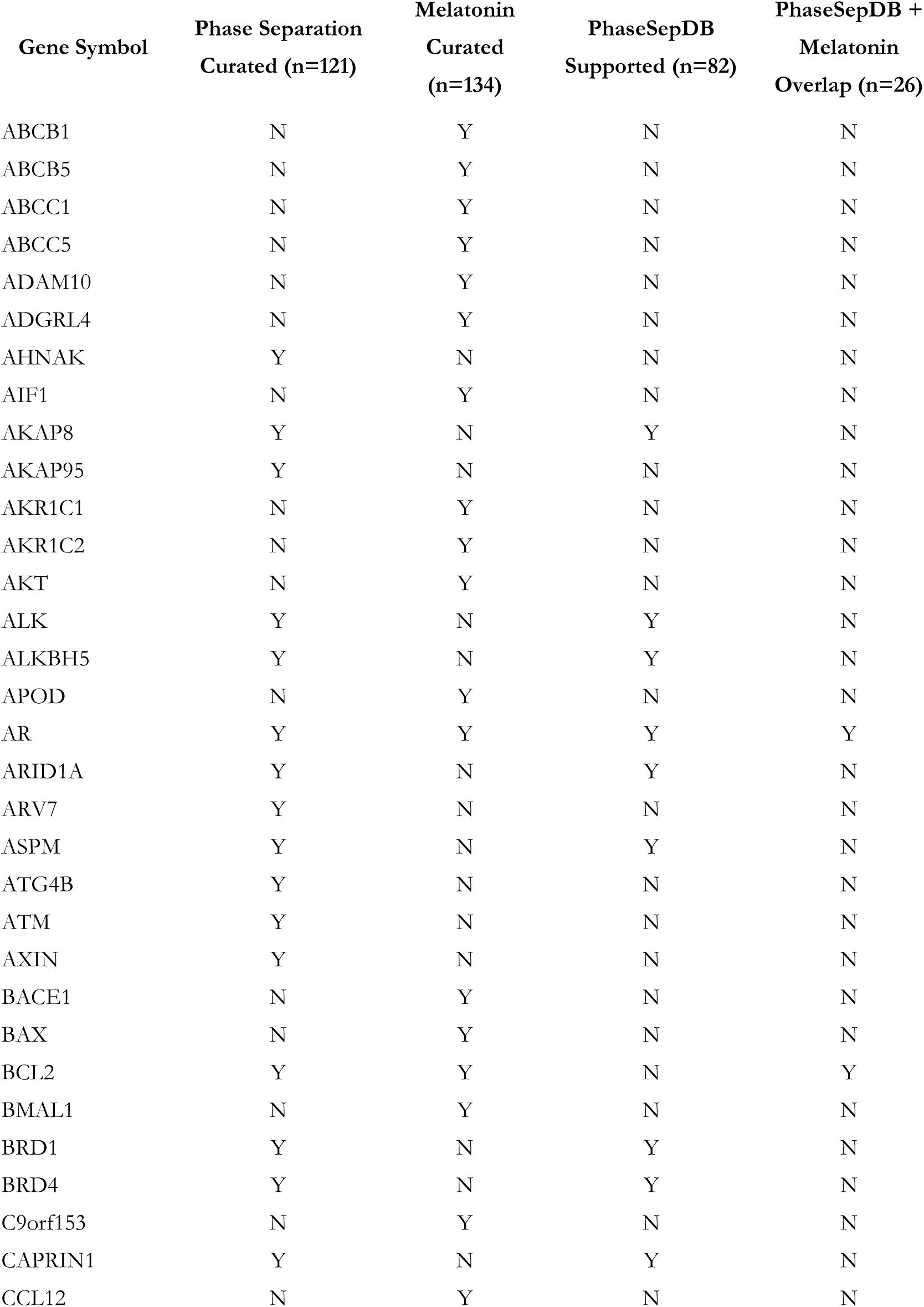
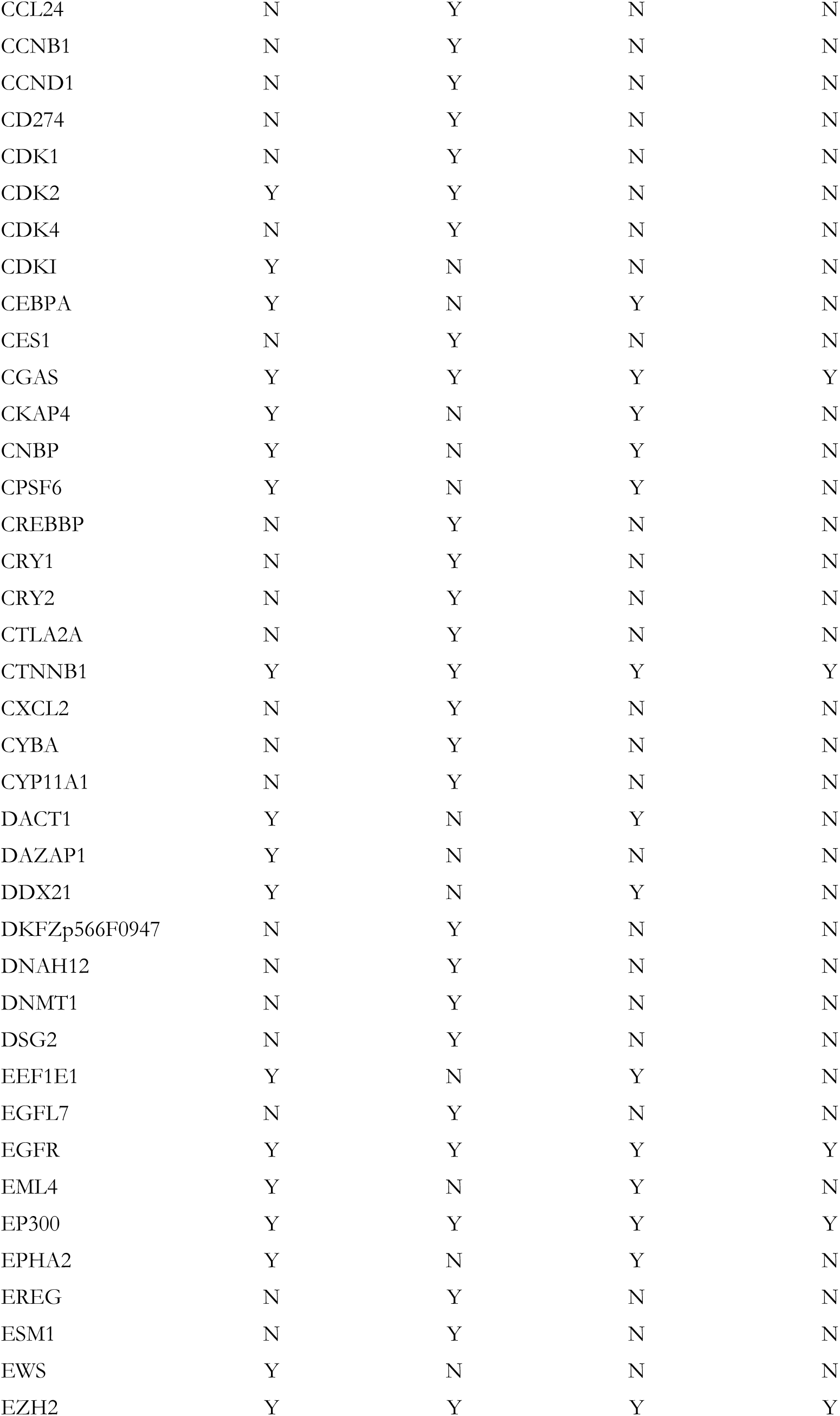
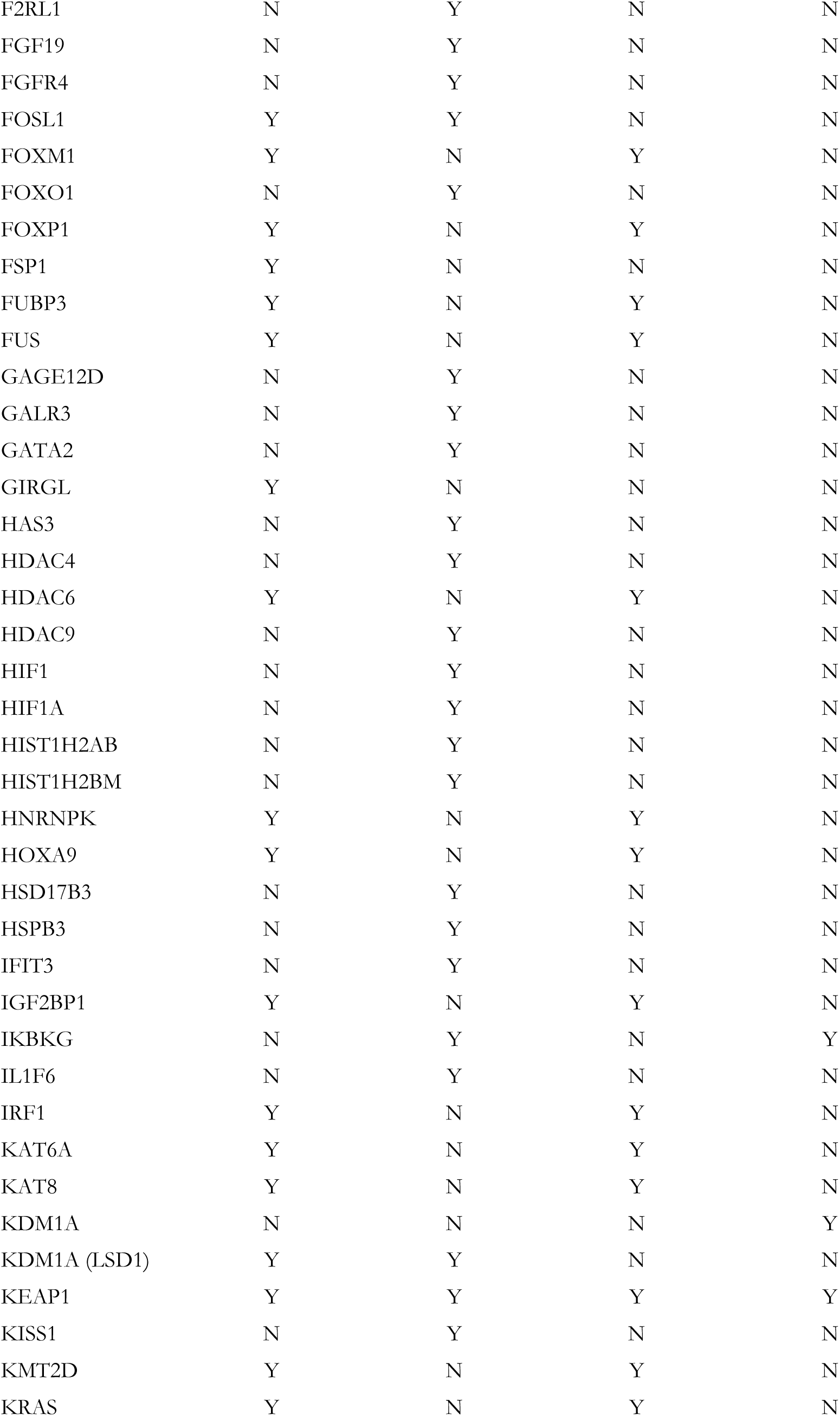
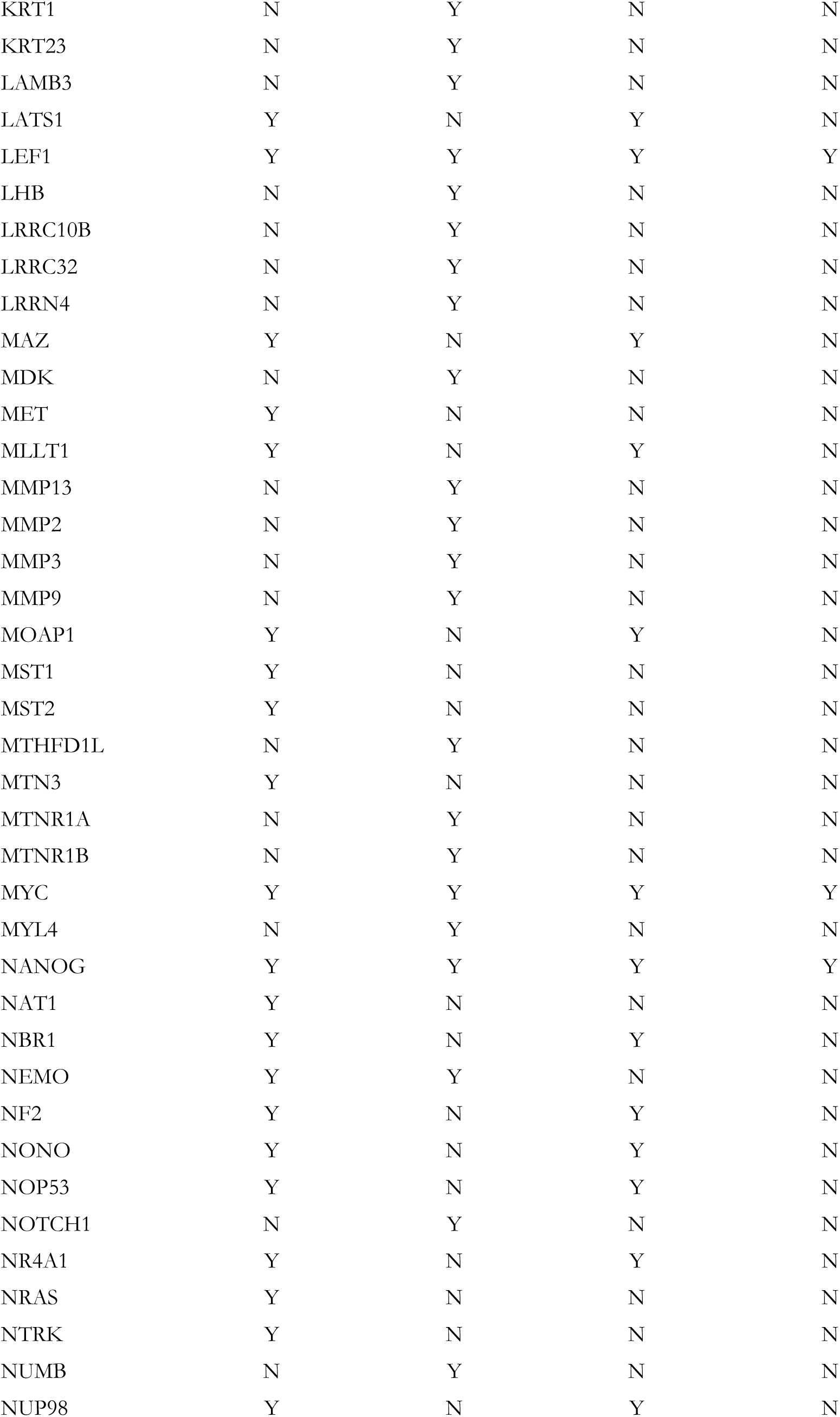
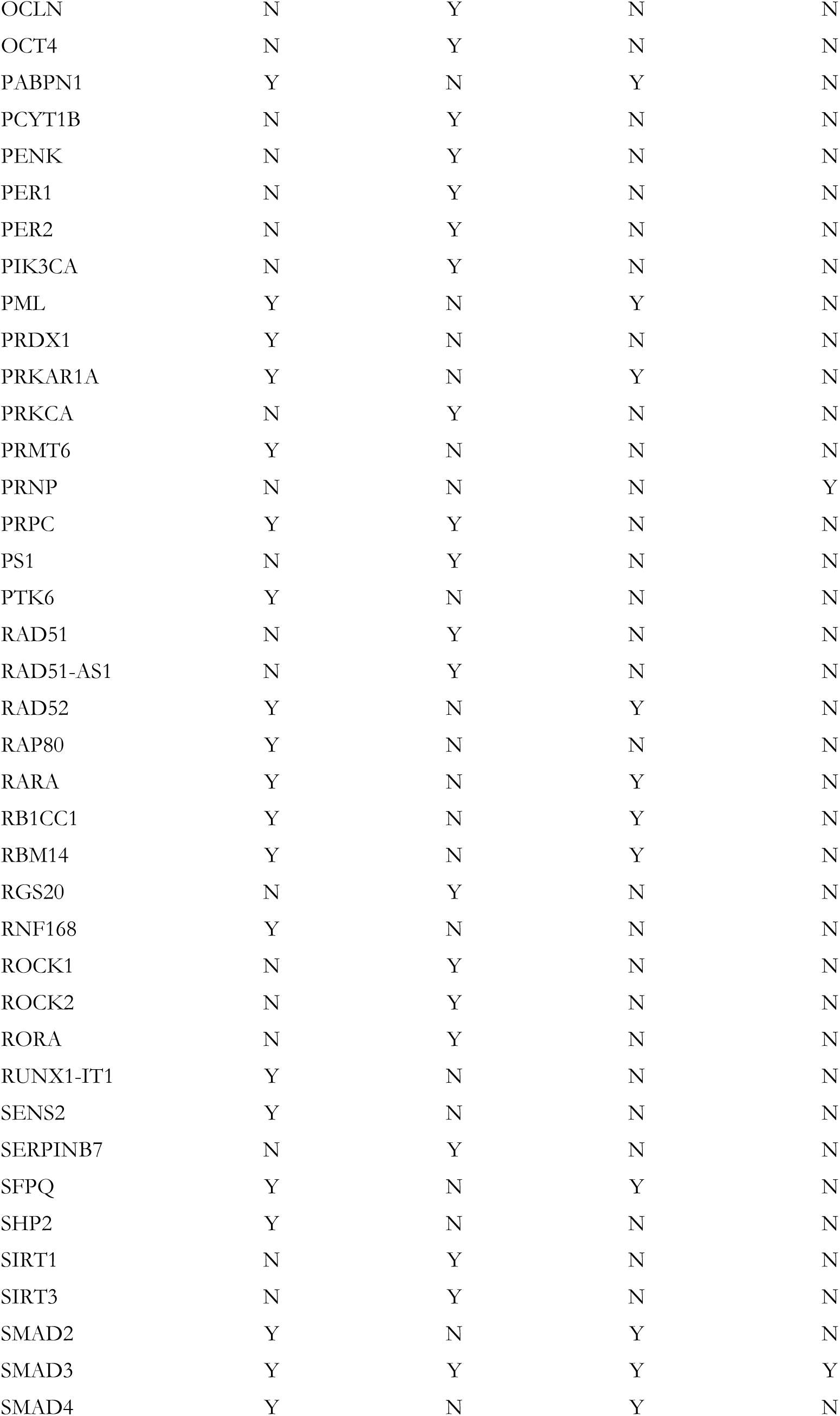
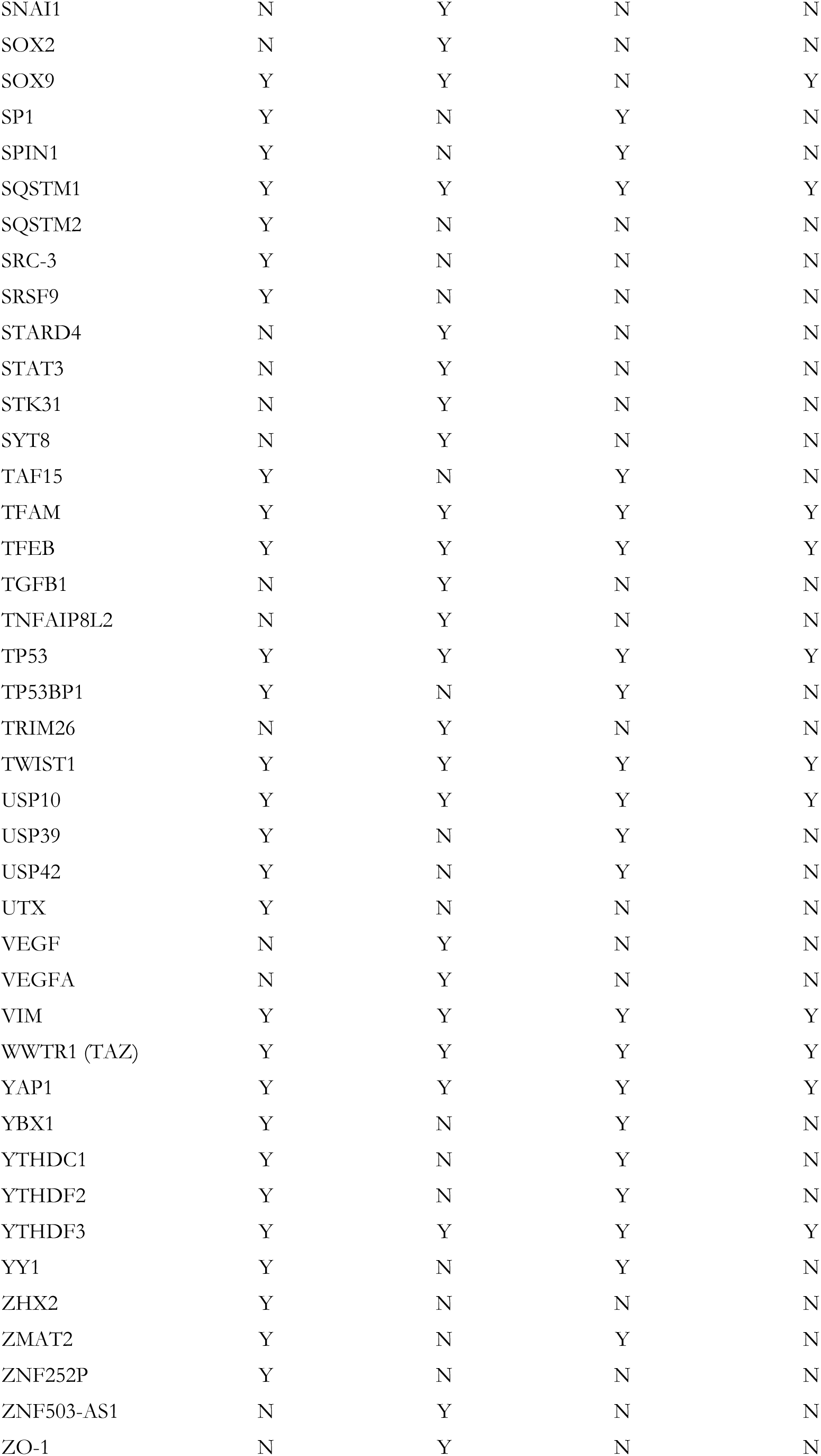
Integrated Bioinformatic Synthesis: Overlap analysis of phase separation, melatonin, and PhaSepDB-validated genes. \*\*\**Table 1 near here – Note: To facilitate a seamless review of the theoretical framework, Table 1 has been placed at the end of the manuscript (after the References) due to its length. This is a primary data element, and we are happy to reposition it per typesetting requirements upon acceptance*..*** *‘Y’ indicates the gene is present in the specified dataset; ‘N’ indicates it was not identified in that specific category. Overlap (n=26) denotes genes validated by PhaSepDB that are also modulated by phase separation and melatonin.

From studies investigating melatonin’s actions across different cancer types, we identified 134 genes significantly modulated by melatonin treatment. Among them, 38 genes were upregulated in at least ten cancer models, predominantly linked to apoptosis, the p53 pathway, Wnt/β-catenin signaling, and TNF-α signaling (MSigDB; FDR < 0.05). In contrast, 105 genes were downregulated, and these were associated with apoptosis, the G2-M checkpoint, KRAS signaling, PI3K/AKT/mTOR signaling, apical junctions, hypoxia, EMT transition, and glycolysis (MSIGDB; FDR < 0.05) (Table 1). Network analysis revealed 22 genes with high-confidence molecular interactions (score = 0.9) (Figure 3A). These genes were mostly localized to the nucleoplasm, nucleus, organelle lumen, membrane-bounded and non-membrane bounded organelles, cytosol, chromosome, survivin complex, extracellular vesicles, among others. Importantly, with the exception of the overexpressed *TP53* and *IKBKG* genes, melatonin significantly downregulated several oncogenic or regulatory genes, including *SOX9, SMAD3, TWIST1, EP300, VIM, LEF1, KDM1A, CTNNB1, EZH2, MYC, AR, BCL2, NANOG, EGFR, USP10, CGAS, YTHDF3, KEAP1, WWTR1, PRNP,* and *TFEB* (Figure 3B).

**Figure 3:**
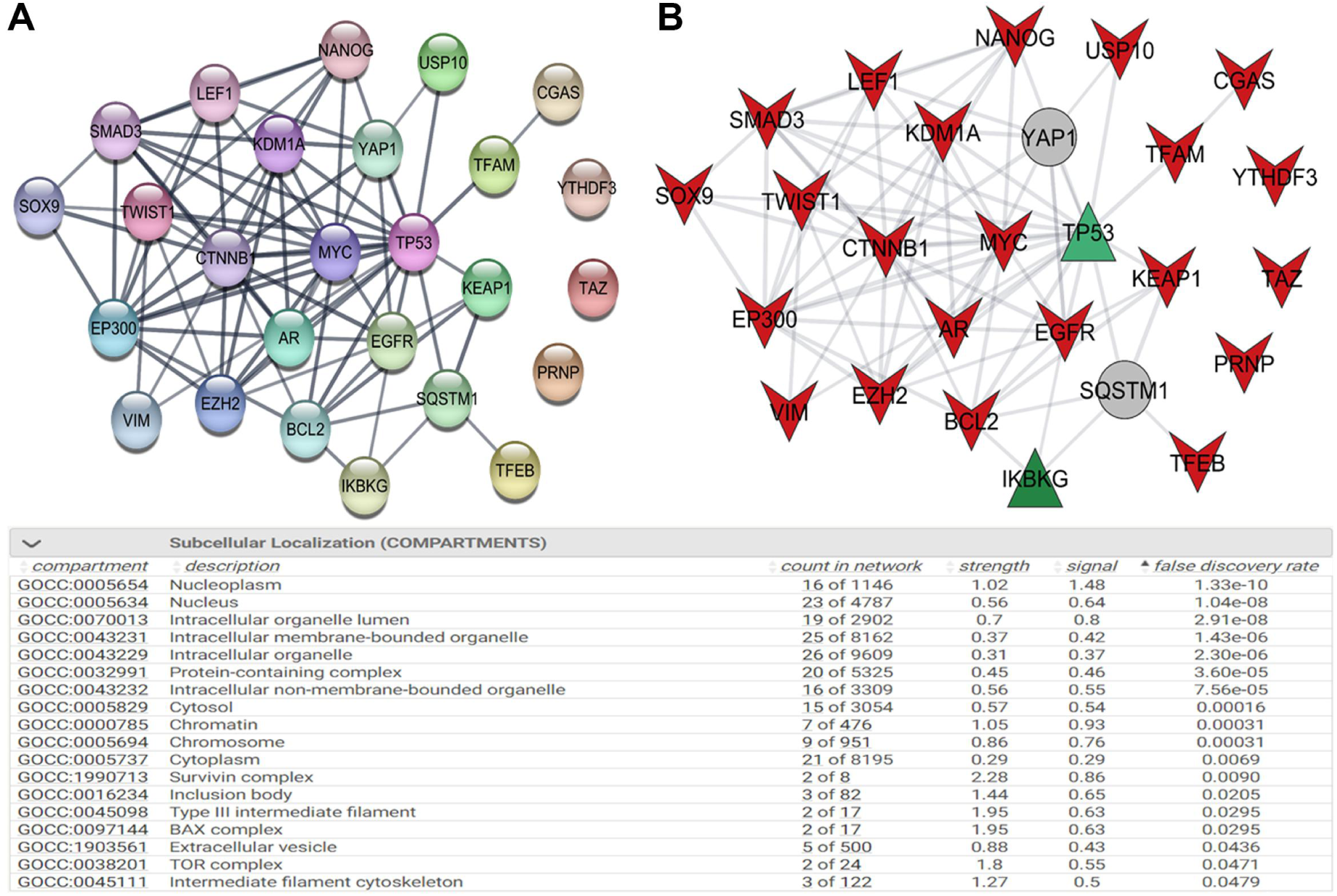
Interaction networks and subcellular enrichment analysis of phase-separation– and melatonin-related genes in cancer. A) STRING-derived interaction network showing the connectivity among selected genes involved in phase separation and responsive to melatonin. Node colors represent distinct functional clusters, and edge thickness indicates interaction confidence. B) Network highlighting common genes and its expression direction. Red and green nodes indicate respectively downregulation and upregulation by melatonin, and gray nodes denote absence of expression modulation. Lower panel: Enrichment table listing the most significantly overrepresented subcellular compartments among the gene set, including nucleoplasm, cytosol, chromatin, and intracellular organelles, with counts, enrichment strength, signal values, and false discovery rates.

### 3.2 Integration of Melatonin-Responsive and Phase-Separation Genes Reveals Key Regulatory Networks in Cancer

By independently analyzing molecules involved in phase separation and those modulated by melatonin, we identified genes at the intersection of both processes to uncover potential functional links in cancer. A total of 26 genes — including *AR, BCL2, CGAS, EGFR, KEAP1, EZH2, LSD1, YAP1, SQSTM1, SMAD3, LEF1, MYC, NEMO, TP53, PRNP, SOX9, WWTR1 (TAZ), NANOG, TFAM, TFEB, TWIST1, EP300, USP10, VIM, YTHDF3*, and *CTNNB1* — were shared between the two datasets, with most being downregulated by melatonin in cancer. We further mapped these genes to their corresponding transcription factors (TFs), cellular effects, and associated cancer types. As shown in Figure 4A, 18 TFs were directly linked to these targets. Notably, *MYC*-, *TP53*-, *BCL2*-, *EGFR*-, and *VIM*-related regulatory networks were prominently represented, connecting to antitumor effects, apoptosis, proliferation, chemoresistance, and metastatic processes in breast, gastric, and liver cancers, as well as glioblastoma.

**Figure 4:**
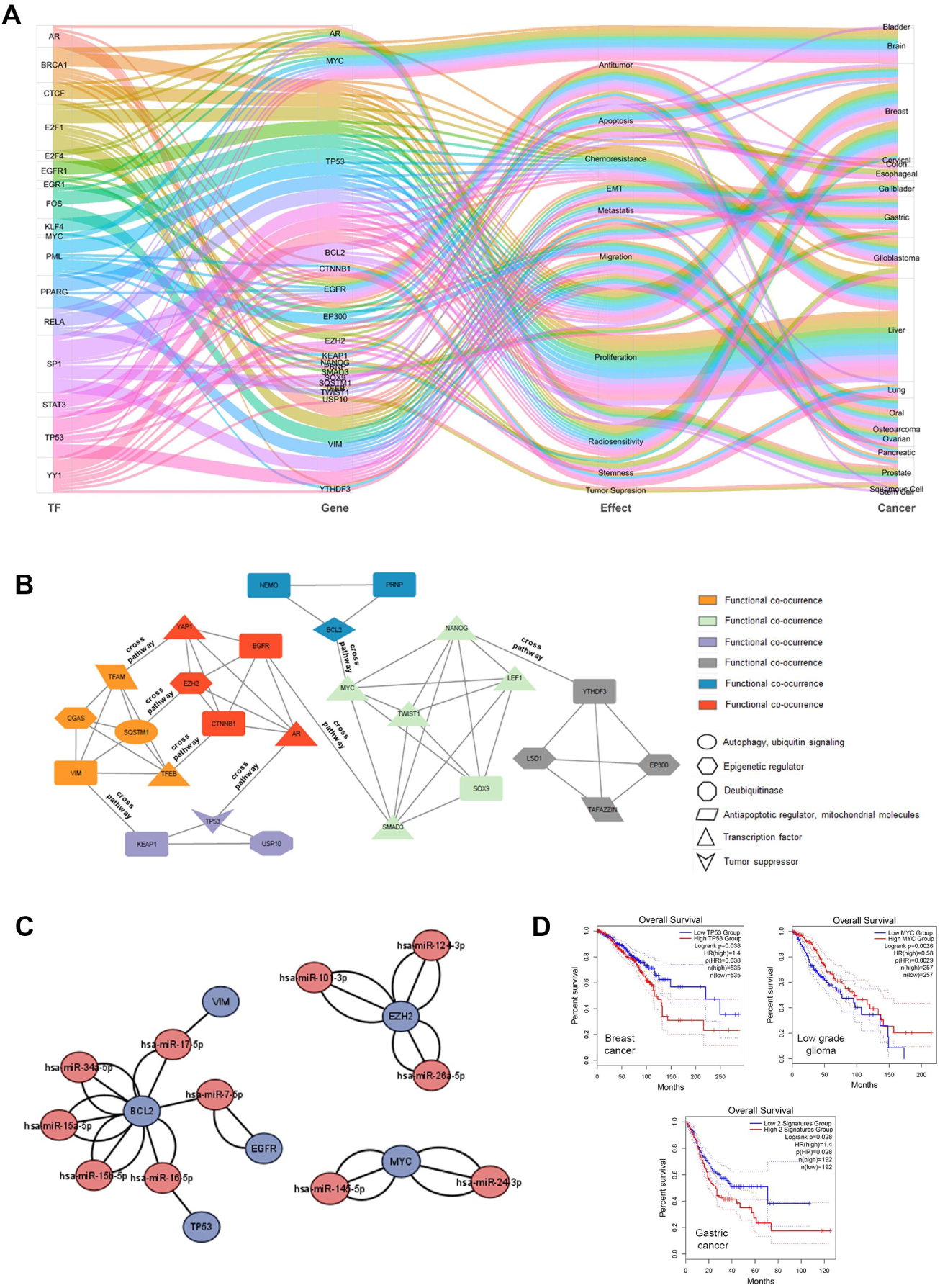
Integrated functional landscape of key regulatory networks in cancer. A) Sankey diagram illustrating the connections between transcription factors (TFs), target genes, their functional effects, and associated cancer types. The visualization highlights major regulatory axes, including TF–gene interactions linked to apoptosis, proliferation, migration, EMT, metabolism, and radioresistance across multiple tumor types such as breast, lung, colorectal, liver, and glioblastoma. B) Functional co-occurrence network showing molecular interactions grouped by biological categories, including autophagy/ubiquitin signaling (yellow), epigenetic regulators (orange), deubiquitinases (blue), autophagic/mitochondrial molecules (green), transcription factors (red), and tumor suppressors (gray). Node clustering emphasizes cooperative functional modules participating in tumor progression. C) MicroRNA–gene regulatory networks showing miRNAs predicted to target core cancer-related genes. Red nodes represent miRNAs and blue nodes represent genes affected by melatonin. D) Kaplan–Meier survival analyses evaluating the prognostic significance of selected gene signatures across different cancer types.

We next clustered melatonin-regulated phase separation genes based on functional co-occurrence, revealing their involvement across interconnected pathways (Figure 4 B). Among these, *LSD1, EP300, CGAS*, and *EZH2* genes enriched in epigenetic regulation, while *TFAM* was associated with mitochondrial functions and *WWTR1* (TAZ) with Hippo-mediated transcriptional regulation. To examine microRNA–gene regulatory interactions, we integrated predictions from three databases, identifying miRNAs targeting core cancer-related genes suppressed by melatonin. For instance, miR-34a-5p, miR-15a-5p, miR-15b-5p uniquely targeted *BCL2* gene; miR-16-5p targeted both *BCL2* and *TP53* genes; miR-7-5p targeted *EGF*R and *BCL2* genes; and miR-17-5p targeted VIM and BCL2. In addition, EZH2 gene was linked to several miRNAs (miR-101-3p, −26a-5p, and −124-3p) whereas *MYC* was associated with miR-145-5p and miR-24-3p (Figure 4C).

To assess clinical significance, we evaluated overall survival of patients using TCGA datasets for high and low risks considering melatonin- and phase separation-related genes. Higher expression levels of specific genes correlated with poorer survival in breast cancer (p=0.038, HR=1.4) and gastric cancer (p=0.028, HR=1.4), whereas low MYC expression was associated with reduced survival rate in low-grade glioma, underscoring the prognostic relevance of these regulatory genes.

## 4 Discussion

Recent advances in the study of phase separation have revealed fundamental organizing principles of biological systems. This rapidly expanding literature has developed largely independent of melatonin [65–67]. Here, we present a synthesis of the extensive evidence documenting the pleiotropic anticancer effects of melatonin on an integrative framework in which phase separation provides a physical basis for melatonin-regulated cancer networks.

To facilitate the interpretation of our results, the 26 identified genes are organized into three functional phase-separation axes that reflect predominant cellular roles rather than exclusive molecular functions. Each gene is assigned to a single phase-separation axis based on its primary condensate-associated role. Importantly, all axes assignments reflect the primary organizational logic of the condensate, not the full spectrum of downstream cellular outcomes. Notably, miRNA regulation converged on all three axes, preferentially targeting *MYC, EZH2, BCL2, EGFR, TP53*, and *VIM*—genes that anchor phase-separated hubs across nuclear, signaling, and stress-response condensates.

### 4.1 Nuclear Decision-Making Condensates (Axis I)

Nuclear decision-making condensates facilitate genome organization and gene-expression regulation by concentrating specific proteins and nucleic acids within biomolecular condensates. These condensates function as nuclear “decision-making” hubs that govern fundamental cellular choices—such as proliferation, differentiation, and cell-fate commitment—by dynamically enhancing or suppressing transcriptional programs. The assembly and dissolution of these condensates are tightly coupled to key cellular decisions during development and in response to environmental or genotoxic cues [68,69].

All genes assigned to Axis I—including *MYC, TP53, AR, KDM1A (LSD1), NANOG, SOX9, EP300, EZH2, LEF1, SMAD3,* and *YTHDF3*—contain intrinsically disordered regions (IDRs) that enable the formation of phase-separated condensate hubs within the nucleus. These hubs often localize to specific genomic regions, such as super-enhancers, where they coordinate transcriptional output. Within the nucleus, proteins such as *EZH2, LEF1, SMAD3*, and *YTHDF3* are frequently recruited into Axis I condensates as regulatory or signaling intermediates. This recruitment enables the integration of upstream signaling inputs into transcriptional outcomes that determine cellular responses, including DNA repair or programmed cell death [13,70–73].

### 4.2 Signal Integration and State Transition Condensates (Axis II)

Signal integration and state transition condensates comprise assemblies that integrate mechanical, biochemical, and spatial signals to drive discrete transitions between cellular states. Canonical examples include YAP/TAZ-, CTNNB1-, and EGFR-associated condensates, which act as molecular decision hubs that convert graded extracellular and intracellular inputs into switch-like transcriptional and phenotypic programs such as proliferation, differentiation, epithelial–mesenchymal transition (EMT), or quiescence [5,74]. Structural and transcriptional modulators such as VIM and TWIST1 have been reported to couple cytoskeletal mechanics and lineage-defining transcriptional programs to condensate-associated signaling thresholds, thereby reinforcing commitment to transitional cell states. VIM can assemble into dynamic non-filamentous states, including biomolecular condensates, enabling rapid reconfiguration in response to mechanical and biochemical cues [75]. In cancer, these adaptable VIM assemblies contribute to epithelial–mesenchymal transition, invasion, and metabolic rewiring, reinforcing commitment to transitional cellular states [76]. Notably, TWIST1 forms phase-separated condensates with YY1 and p300 at super-enhancers to drive oncogenic transcriptional reprogramming in hepatocellular carcinoma [77]. In parallel, IKBKG (NEMO)-containing nuclear signaling condensates integrate inflammatory and stress-associated cues to bias NF-κB–dependent transcription toward state reprogramming outcomes [78]. Although these condensates can secondarily promote stress tolerance through transcriptional rewiring, their defining feature is the orchestration of context-dependent state transitions rather than the direct buffering of cellular stress.

Whereas Axis II condensates mediate the decision to enter a particular cellular state, Axis III condensates stabilize survival within that state under adverse conditions. Notably, YAP/TAZ can also participate in stress-induced condensates that enhance cellular persistence under mechanical or metabolic constraint; such assemblies are classified here under Axis III when their dominant function shifts from state transition to stress endurance

### 4.3 Stress Adaptation and Survival Condensates (Axis III)

Stress adaptation and survival condensates are distinguished by their switch-like sensitivity to environmental stress, enabling rapid transitions between cellular preservation and collapse. Axis III condensates primarily function as reactive storage and protective hubs that buffer oxidative, metabolic, and proteotoxic stress [79]. In contrast to Axis I condensates, which support continuous information flow and transcriptional regulation, Axis III condensates prioritize immediate cell survival, determining whether cancer cells tolerate stress or undergo irreversible collapse.

Central to this axis are stress-responsive condensates organized around USP10, SQSTM1 (p62), and stress granules (SGs) that sequester signaling components to bias cellular programs toward survival While transient formation of SGs is protective under acute stress, their persistent stabilization and accumulation under chronic stress conditions—such as those encountered in tumors—can promote pathological survival states [80]. A defining feature of Axis III regulation is the sequestration of KEAP1 into SQSTM1-containing condensates, enabling NRF2-dependent cytoprotective gene expression and antioxidant adaptation [81]. In addition to buffering metabolic and proteotoxic stress, Axis III condensates modulate immune and apoptotic thresholds. Phase separation of cGAS serves as a switch leveraged by cancer cells to drive metastasis (on) or evade immune detection (off) [82]; while condensate-associated interactions involving PrP^c^ (*PRNP*) and BCL2 raise apoptotic thresholds under stress [83]. In parallel, phase separation–linked regulation of TFEB and TFAM supports lysosomal and mitochondrial programs that reinforce metabolic flexibility and stress tolerance [84–86]. Collectively, Axis III condensates function as stress-responsive decision nodes, enabling cancer cells to survive hostile microenvironments through dynamic molecular buffering and survival prioritization

The assembly of phase-separated condensates in all three axes are evolutionarily conserved, transient, physiological responses to acute stress. Under chronic stress conditions, including cancer, phase separation transitions into an aberrant phase, becoming drivers and promoters of oncogenesis [87]. The tumor microenvironment can modulate biophysical parameters that affect condensate behavior, leading to aberrant phase separation. Condensates in the three phase separation axes are extremely sensitive to several biophysical axes of control that regi;ate condensate behavior [88]. Accordingly, the 26 genes all exhibit sensitivity to multiple biophysical levers, not surprisingly, known to be modulated by melatonin. For clarity, we refer to these biophysical axes as control levers when discussing their modulatory effects on genes and condensates.

### 4.4 Melatonin Modulates Condensate Biophysical Levers

An important conceptual consideration in interpreting melatonin-regulated gene expression in cancer studies is that bulk quantification approaches, which dominate the cancer literature, report net transcript or protein abundance but are inherently agnostic to mesoscale organization of molecules assembled via phase separation. Consequently, observed melatonin-induced up- or downregulation of phase separation–associated proteins does not preclude their simultaneous partitioning into biomolecular condensates, nor does it exclude melatonin-mediated modulation of the phase separation process itself. While bulk measurements capture changes in total transcript or protein abundance, phase separation has been increasingly implicated in modulating gene expression through the formation of transcriptional condensates that concentrate or sequester regulatory factors and thus influence transcriptional outputs in a dynamic, context-dependent manner [68,89]. In this context, changes in bulk expression may reflect downstream consequences of altered condensate formation, stability, or material properties, as well as feedback regulation arising from condensate-dependent transcriptional or post-transcriptional control.

The identification of 26 genes jointly associated with phase separation and melatonin regulation, therefore, supports a model in which melatonin influences not only protein abundance but also the biophysical state and functional compartmentalization of key cancer-relevant regulators—effects that are not resolvable by bulk measurements alone. Notably, the three phase-separation axes are defined not only by biological function, but also by distinct biophysical sensitivities—redox state, multivalent interaction strength, and electrostatic balance—known to be influenced by melatonin. The existence of a sovereign singularity, where phase-separation architecture and melatonin biology converge to govern health and disease, warrants further elucidation.

#### 4.4.1 Redox Tuning of Condensate Stability (Lever I)

Since the origins of life, redox (reduction-oxidation) chemistry and phase separation have been intrinsically linked in cellular organization and stress response mechanisms [90]. Redox-controlled phase separation acts as a molecular switch that enables cells to sense, respond, and adapt to environmental and metabolic fluctuations by governing the rapid formation and dissolution of BCs [91–93]. In the pursuit of survival and proliferation in a hostile environment of persistent oxidative stress and a reversed pH gradient, cancer cells rebalances cellular redox homeostasis [94], upregulating antioxidant defenses by flipping protein redox switches to cause the aberrant phase separation of oncogenic condensates.

This ‘redox-first’ assembly initiates the formation of functional hubs across the cell’s architecture: from nuclear decision-making (Axis I) driven by LSD1, EP300, MYC, and EZH2, to the signal integration (Axis II) of EGFR, and the stress adaptation (Axis III) of KEAP1 and BCL2. While redox reactions trigger the initial molecular switch—often mediated by redox-sensitive post-translational modifications [95]—the ultimate stability and material properties of these condensates are dictated by an intricate interplay with multivalent scaffolding (Lever II) and electrostatic control (Lever III).

#### 4.4.2 Modulation of Multivalent interactions (Lever II)

While redox reactions flip the initial molecular switch, the structural persistence and material properties of oncogenic condensates are maintained by multivalent interactions. Lever II governs the cooperative cross-linking of the dense phase, utilizing a diverse chemical toolkit—including hydrophobic effects, cation-π interactions, π-π stacking, and hydrogen bonding—to create interconnected molecular networks [96]. Notably, while many condensates rely on intrinsically disordered regions (IDRs), multivalent scaffolding in several key oncogenic drivers occurs independently of classic disorder. For instance, the structural integrity of CTNNB1 (β-catenin) and SQSTM1 is mediated primarily through folded domains and repetitive binding interfaces rather than disordered motifs [97,98].

In the cancer TME, Lever II is exploited to stabilize nuclear decision-making (Axis I) through the recruitment of AR, LEF1, SMAD3, and SOX9, forming high-density transcriptional hubs. Similarly, signal integration (Axis II) is amplified via the multivalent assembly of the Wnt and Hippo pathways—specifically through CTNNB1, TAZ (*WWTR1)*, YAP1, and TWIST1. Furthermore, in stress adaptation (Axis III), proteins like PrP^c^ (*PRNP*) and SQSTM1 leverage their domain-based multivalency to sequester vital cellular components, providing a physical shield against apoptotic signals [99]. This layer of structural reinforcement ensures that these aberrant condensates remain resilient to transient environmental fluctuations, representing a critical therapeutic vulnerability in tumorigenesis [5].

#### 4.4.3 Electrostatic and Stoichiometric Control (Lever III)

While multivalency provides the structural scaffold, electrostatic and stoichiometric control dictate the selectivity and internal environment of the dense phase of condensates. Lever III governs the partitioning of ions and the net charge-density matching within the condensate, creating a localized bio-electrochemical field that regulates cellular electrochemical equilibria [100]. Through a process of charge neutralization, these hubs can sustain significant pH gradients at equilibrium, effectively decoupling the internal p*K*_a_ from the increasingly acidic TME [101]. In the cancer TME, the reversed pH gradient acts as a biophysical catalyst for these interactions, promoting a Proton Trap mechanism. Recent all-atom continuous constant pH molecular dynamics simulations demonstrate that condensate microenvironments universally favor protonated states by acting as unique solvation environments that reshape the p*K*_a_ values of amino acid side chains. This shift stabilizes the charged forms of cationic residues (HIS, LYS) and the neutral forms of anionic residues (ASP, GLU), fundamentally altering the solvation properties that govern biochemical function [102,103].

These physicochemical shifts drive aberrant phase transitions, now recognized as an emerging hallmark of cancer [5]. While metabolites universally shift the thermodynamic equilibria of biomolecular condensates[104], the unique metabolic profile of the cancer TME—characterized by high interstitial fluid pressure and a surplus of molecular crowders and nucleic acids—exploits this physical sensitivity. These conditions drive the aberrant phase transitions now recognized as a hallmark of cancer, effectively strengthening the electrostatic forces that stabilize oncogenic hubs. This electrostatic “shield” is exploited across all functional axes: nuclear regulators (Axis I) such as NANOG, EZH2, and YTHDF3 utilize charge-density matching to lock in transcriptional programs [105], while signal integration (Axis II) hubs like EGFR, IKBKG (NEMO), and VIM leverage increased positive charge to amplify pro-survival signaling [78]. Most critically, for stress adaptation (Axis III), the formation of cGAS, TFEB, USP10, and BCL2 condensates is stabilized by this charge-dependent insulation, as exemplified by TFAM, which protects mitochondrial genomic integrity [106,107]. By exploiting the Proton Trap, cancer cells ensure that oncogenic survival programs remain biophysically viable despite the hostile extracellular landscape.

### 4.5 Melatonin: The Sovereign Singularity of the Field Effect in Biomolecular Condensate

The complexity of the 26-gene oncogenic network—spanning nuclear hubs (Axis I), signaling cascades (Axis II), and survival adaptations (Axis III)—suggests that current clinical strategies must evolve to match the scale of cancer’s biophysical adaptability. Modern oncology has made significant strides with a ‘combination lock’ approach, utilizing synergistic pairings such as Antibody-Drug Conjugates (ADCs) and Immune Checkpoint Inhibitors (ICIs) to address multiple, distinct facets of tumor biology simultaneously [108]. However, as these therapies target increasingly comprehensive downstream effects, the question arises whether even the most sophisticated designs are perpetually chasing an evolving oncogenic landscape. We propose that melatonin functions as a sovereign singularity, offering a higher-order intervention that transcends the need to pick individual molecular tumblers. By regulating the integrated biophysical parameters that serve as the foundational ‘field’ for these 26 genes, melatonin renders the oncogenic program biophysically unviable, effectively stabilizing the landscape before the next evolutionary evasion can occur.

While the ‘sovereign singularity’ remains a theoretical synthesis, it is grounded in emerging experimental evidence suggesting that melatonin does not merely inhibit proteins, but fundamentally alters the physical state of the oncogenic niche [29,41,58]. By bridging the gap between traditional pharmacology and systems biophysics, our proposed framework offers a strategic roadmap for future experimental validation of melatonin as an evolutionarily conserved, universal regulator of cellular phase behavior (Figure 5).

**Figure 5:**
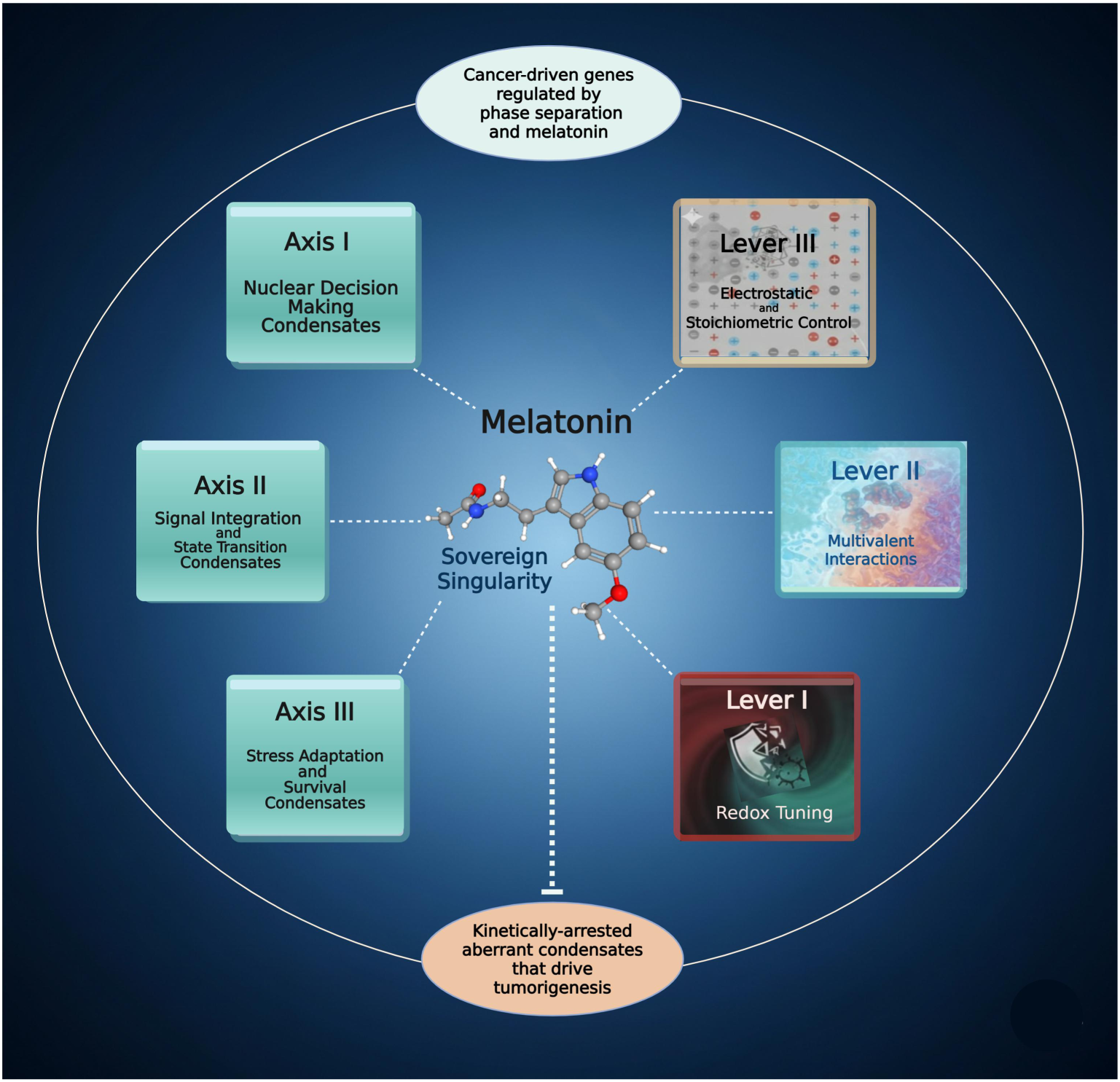
Melatonin: A Sovereign Singularity for Disrupting Oncogenic Phase-Separated Networks by targeting integrated biophysical axes (Axes 1–3) that stabilize oncogenic phase-separated condensates. By modulating a tri-lever framework (Levers 1–3), melatonin recalibrates redox homeostasis, disrupts multivalent interactions acting as a molecular plasticizer, and normalizes electrochemical energetics by functioning as a dielectric tuner. By exerting a field effect in the tumor environment, melatonin enforces fundamental physicochemical constraints that render the oncogenic survival program biophysically unviable.

#### 4.5.1 Melatonin Recalibrates Redox Homeostasis as the Ultimate Phase-Transition Switch (Lever I)

The traditional view of melatonin as a monolithic antioxidant is increasingly challenged by its context-dependent “pro-oxidant” effects in malignant cells. Recent evidence demonstrates that melatonin can paradoxically reduce glutathione (GSH) and catalase (CAT) levels in ovarian cancer, promoting cytotoxicity while stripping the cell of its oxidative defenses [109][. Within the sovereign singularity framework, this effect is not a contradiction but a strategic biophysical recalibration. In the TME, cancer cells utilize redox-driven phase separation to sequester and stabilize antioxidant machinery—such as the NRF2/KEAP1 and MCM5 hubs—effectively creating a ‘redox shield’ that permits survival under high oxidative stress [58]. The paradoxical pro-oxidant and cytotoxic effects of melatonin represent a systemic phase reset rather than mere chemical reactions. When melatonin terminates the aberrant thermodynamic conditions that allow antioxidant enzymes and metabolites (CAT/GSH) to be sequestered and hyper-stabilized within protective condensates, it effectively strips the cancer cell of its biophysical armor, exposing vulnerabilities that lead to the demise of the malignant cell. This recalibration ensures that the redox landscape is returned to physiological baseline, rendering the 26-gene survival network biophysically exposed.

#### 4.5.2 Melatonin Disrupts Multivalent Scaffolding as a Molecular Plasticizer (Lever II)

While the redox switch initiates the transition, the persistence of oncogenic hubs depends on the multivalent ‘stickiness’ of intrinsically disordered regions (IDRs) of proteins in Lever II. Proteins such as CTNNB1 (β-catenin) and YAP1 rely on transient π-π and cation-π interactions to form dense, gel-like scaffolds. We propose that melatonin functions as a molecular plasticizer that disrupts the physical sanctuary for pro-survival signaling resistant to standard enzymatic degradation. The electron-rich indole ring and dipole moment allow melatonin to partition into proteins with IDRs, competing for the aromatic contacts that drive cross-linking [57,110–112]. By disrupting this multivalent molecular grammar, melatonin maintains the fluidity of the 26-gene assembly, preventing it from maturing into a rigid, irreversible signaling haven As a molecular plasticizer, melatonin ensures that oncogenic scaffolds remain dynamic and accessible to the physiological degradation machinery.

#### 4.5.3 Melatonin Normalizes Electrochemical Energetics as a Dielectric Tuner (Lever III)

To complete the tri-lever triad, we move from the internal architecture of the condensate to its relationship with the surrounding environment. This is where the sovereign role of melatonin is most apparent, as it governs the very energetics of the cellular solvation and fluidity landscape. Cancer cells exploit the Proton Trap in Lever III in order to maintain aberrant condensates with internal p*K*_a_ values that are decoupled from the increasingly acidic TME. Recent evidence confirms that such condensates can sustain significant pH gradients relative to the bulk environment at equilibrium through a process of charge neutralization [101]. The decoupling of condensate internal p*K*_a_ from the increasingly acidic TME provides a biophysical sanctuary that renders Axis III survival programs resistant to extracellular stress [102].

Melatonin collapses cancer cell Lever III defenses through a dual mechanism of metabolic recalibration and direct dielectric tuning. By shifting the cellular program from glycolytic fermentation to oxidative phosphorylation, melatonin reduces the accumulation of metabolic “crowders” that shift thermodynamic equilibria [104,113]. Simultaneously, melatonin—particularly when stacked with ATP—modulates the local dielectric constant (ε) within the condensate environment [57]. The synergistic stacking between the adenine purine ring and the melatonin indole ring creates a high-dipole molecular complex that acts as a biophysical buffer, neutralizing the charge-gradient effects identified by Ausserwöger et al. (2025). By dissipating these localized electrochemical fields, melatonin forces the oncogenic hubs to re-equilibrate with the physiological environment, rendering the 26-gene and other potential cancer networks biophysically untenable and accessible to therapeutic intervention [29]. Table 2 summarizes the 26 phase-separation cancer genes regulated by melatonin; it details their expression patterns (upregulation/downregulation), biophysical levers, functional axes, and roles in cancer progression. Please refer to Table to for synonyms and full names of the 26 genes.

**Table 2:**
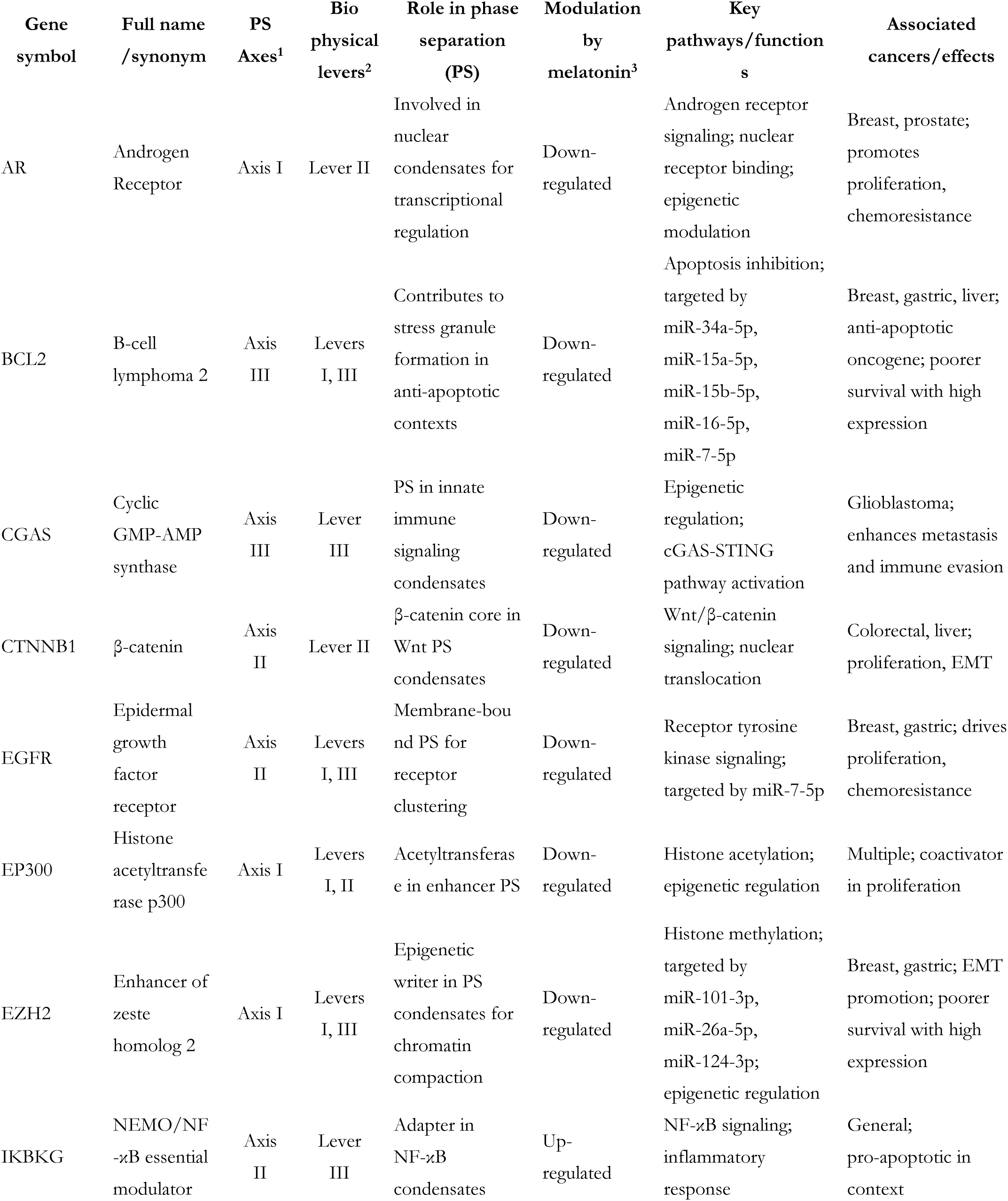
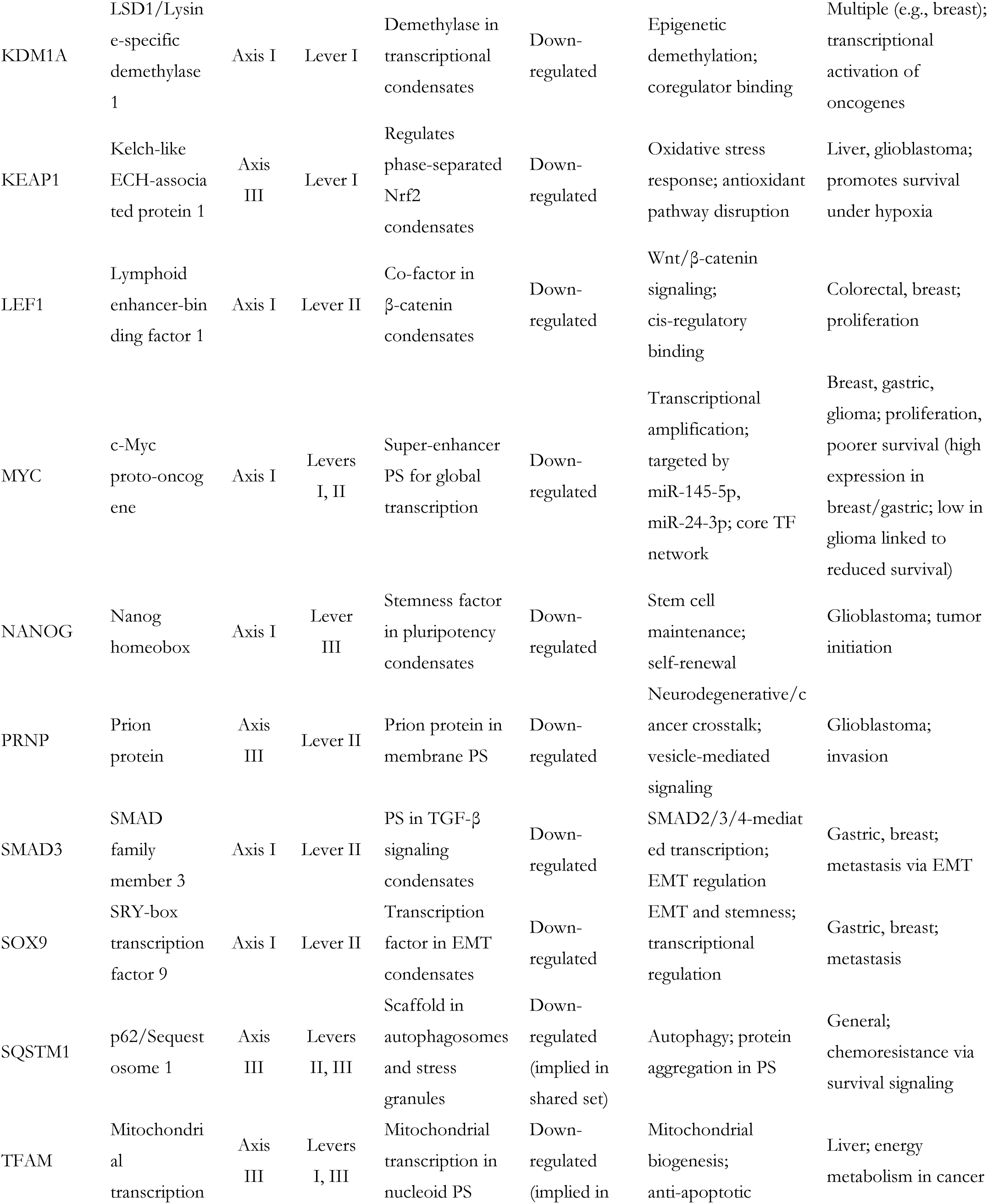
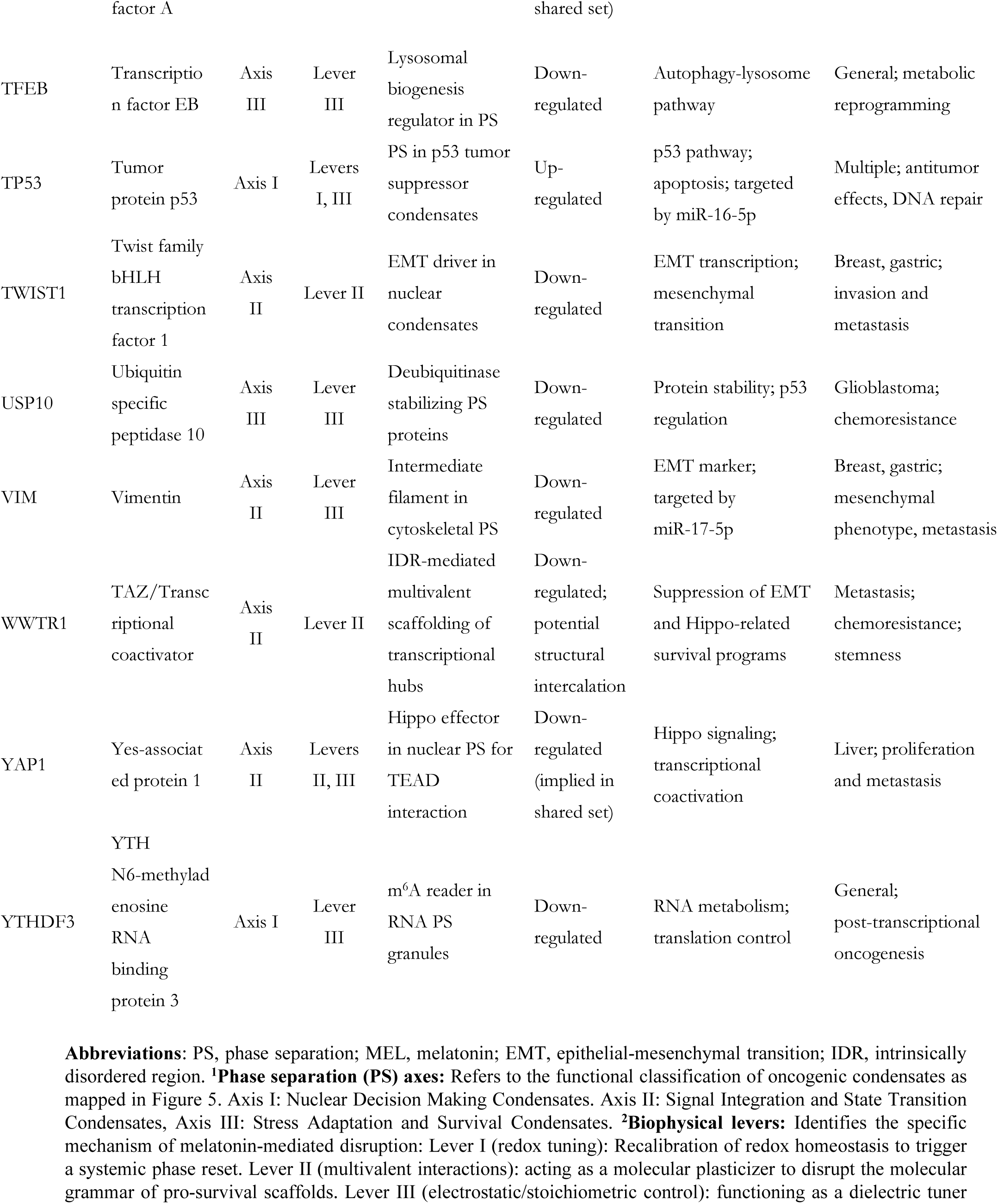
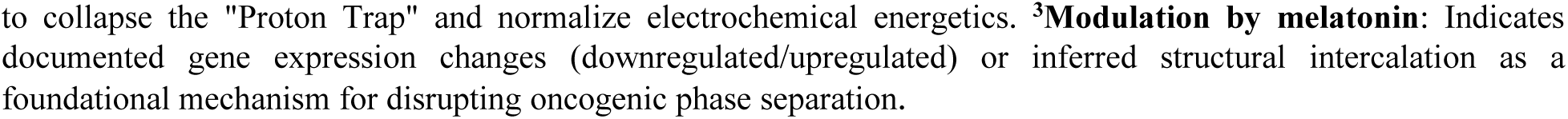
Bioinformatic convergence analysis: Biophysical and functional profiles of genes regulated by melatonin and phase Separation.

## 5 Limitations

While our proposed framework provides a novel unification of melatonin’s pleiotropic effects, several conceptual boundaries must be noted. First, the three biophysical levers identified here are representative of the primary regulatory forces currently known to govern cellular organization. They are intended as a fundamental template for biophysical regulation rather than an exhaustive catalog of all potential force-fields involved in phase behavior. Second, the 26-gene survival network synthesized in this study provides strategic coverage of core oncogenic hubs across multiple functional axes. As the study of biomolecular condensates expands, the list of validated targets will undoubtedly grow. However, the consistent modulation of this representative core suggests that the underlying biophysical principles—the “sovereign” regulation of the cellular environment—remain the universal common denominator of melatonin’s efficacy, regardless of the specific genes involved.

## 6 Conclusions and Future Perspectives

Our integrative bioinformatics proof-of-concept analysis reveals melatonin acting as a universal regulator of cellular fluid dynamics, ensuring biophysical and biochemical mechanisms that drive phase separation—the spontaneous organization of cytoplasmic and nucleoplasmic assemblies—remain physiologically viable and functional. By permeating and destabilizing the biophysical sanctuaries that enable cancer’s evolutionary adaptability, melatonin effectively “de-shields” the malignant cell, rendering its survival machinery biophysically untenable.

This proof-of-concept synthesis raises a fundamental question: is melatonin merely a participant in this biophysical convergence, or is it the sovereign singularity itself? Given its unique capacity to integrate redox tuning, multivalent interaction, and electrostatic balance, melatonin likely serves as the defining coordinate of this singularity—the singular point where the cell’s physical architecture converges with its survival logic. In the search for a universal regulator of life and a singular vulnerability in cancer, melatonin may not simply be a part of the solution; it may be the biophysical constant that defines the equilibrium of the living state.

To move this framework from a provocative hypothesis to a definitive biological principle, future studies must evolve beyond traditional expression profiling. Research integrating transcriptomic and proteomic data with spatially resolved or condensate-sensitive assays will be necessary to discern abundance-based effects from phase state–dependent regulation in melatonin-responsive cancer pathways. Such experimental validation will ideally address the full depth of the biophysical levers and how they modulate phase separation axes. The therapeutic significance of this convergence lies in the transition from molecular targeting to landscape-level biophysical intervention. By modulating the fluid properties of cancer gene networks, melatonin strategically manages the diverse evasion toolkits used by malignant cells to evade inhibition. Furthermore, when deployed at precise physiological or pharmacological molarities, melatonin serves as a critical combinatorial adjuvant; by enforcing fundamental physiological parameters in the biophysical landscape, it ensures that the specific mechanisms of concurrent therapies are optimized and that the survival of cancer cells—critically dependent upon aberrant condensate assemblies—becomes untenable.

## Acknowledgement

Special thanks to Daniel Matrone for technical assistance. Figure 5 created in https://BioRender.com with elements generated by Gemini 3 Flash.

## Author Contributions

The authors confirm contribution to the paper as follows: Doris Loh: conceptualization, methodology (systematic review), writing—original draft, writing—review & editing. Luiz Gustavo de Almeida Chuffa: methodology (bioinformatics), software, formal analysis, writing—original draft, writing—review & editing. Fábio Rodrigues Ferreira Seiva: methodology (bioinformatics), software, formal analysis, writing—original draft, writing—review & editing. Russel J. Reiter: writing—critical review & editing. All authors reviewed the results and approved the final version of the manuscript.

## Availability of Data and Materials

The data that support the findings of this study are available from the Corresponding Author, Doris Loh, upon reasonable request.

## Ethics Approval

Not applicable.

## Supplementary Materials

Not applicable.

## Glossary/Nomenclature/Abbreviations

Abbreviation: Full Term
ADCs: Antibody-Drug Conjugates
BCs: Biomolecular Condensates
CAT: Catalase
EMT: Epithelial–Mesenchymal Transition
GO: Gene Ontology
GSH: Glutathione
ICIs: Immune Checkpoint Inhibitors
IDRs: Intrinsically Disordered Regions
LSD1: Lysine-specific Demethylase 1A (KDM1A)
NEMO: NF-kappa-B Essential Modulator (IKBKG)
PPI: Protein-Protein Interaction
PRISMA 2020: Preferred Reporting Items for Systematic Reviews and Meta-Analyses
PSepDB: Phase Separation Database
TAZ: Transcriptional Coactivator with PDZ-binding Motif (WWTR1)
TF: Transcription Factor
TME: Tumor Microenvironment

